# Cardiaca adipokinetic hormone and hedgehog signaling combine to generate intracellular waves of Ca^++^ in starved *Drosophila melanogaster* fat body

**DOI:** 10.1101/2024.04.05.588282

**Authors:** Min Kang, Anthea Luo, Isabelle Becam, Anne Plessis, Robert A. Holmgren

## Abstract

The *Drosophila melanogaster* fat body combines the functions of the vertebrate liver and fat. It plays a central role in metabolism where it integrates information about nutritional status to regulate fat utilization. During feeding, signaling through the Insulin Receptor causes lipogenesis, while fasting leads to signaling through the cardiaca Adipokinetic Hormone Receptor (AKHR) and mobilization of lipid stores. Here we examine intracellular calcium levels in the fat body during fasting. In fasting early third instar larvae, spikes of intracellular calcium are generated in the fat body lobes on either side of the brain. These spikes propagate through a narrow connection into the main lobes of the fat body that lie along the length of the larva. The spikes of intracellular Ca^++^ are dependent on the corpora cardiaca AKH expressing neurons and AKHR. Unexpectedly, the spikes also require Hedgehog (Hh) signaling from the midgut enterocytes and within the fat body. When Hh signaling is blocked, the Ca^++^ levels in the fat body are elevated and the spiking behavior lost. Hh signaling appears to regulate fat body intracellular Ca^++^ using both the transcription factor Cubitus interruptus and the trimeric G protein Gαi. AKH/Hh signaling in the fat body lobes on either side of the brain appears to function as a pulse generator to initiate Ca^++^ spikes that then propagate through the main lobes of the fat body. These studies show how signaling from the brain and the midgut and within the fat body are integrated to regulate a key intracellular second messenger.

## Introduction

The *Drosophila melanogaster* fat body integrates signals related to nutritional status to regulate metabolism. It carries out the combined functions of the vertebrate liver and adipose tissue. (1-3). In response to fasting, the corpora cardiaca (CC) neurons release Adipokinetic Hormone (AKH) (4-6), the *D. melanogaster* homologue of Glucagon, which binds to the AKH Receptor (AKHR). AKHR signals through Gαq and Gγ1 to activate Phospholipase C and the Inositol 1,4,5 triphosphate Receptor (IP_3_R), leading to elevated intracellular Ca^++^ levels (7-9). In *Manduca sexta*, AKH signaling in the fat body has also been shown to signal through Gαs to activate PKA (10, 11). As a consequence, glycogen and triglyceride stores are released from the fat body (12). Elimination of the AKH producing neurons leads to decreased levels of trehalose (a key *D. melanogaster* sugar) in the hemolymph while overexpression of AKH leads to depletion of lipid droplets in the fat body (5, 13). Fasting also leads to a reduction in insulin signaling, which promotes nuclear translocation of FOXO and the expression of lipases(14, 15).

Hedgehog (Hh) signaling from the midgut enterocytes (EC) has also been implicated in the *D. melanogaster* starvation response (16-18). Loss of Hh signaling from midgut ECs prevents neutral lipid mobilization upon starvation (17). This effect appears to involve direct signaling to the fat body as manipulating the activity of the Hh receptor Patched (Ptc) in the fat body sensitizes the larvae to starvation (17).

In the canonical Hh pathway, binding of Hh to its receptor Ptc blocks Ptc repression of the G protein-coupled receptor (GPCR) Smoothened (Smo), which in turn leads to the activation of the GLI family transcription factors (Cubitus interruptus (Ci) in *D. melanogaster*) (19). In the fat body, starvation activates canonical Hh signaling and the direct transcriptional activation of the *brummer* (*bmm*) lipase gene by Ci (18). While it has been demonstrated that Smo can also couple to Gαi (20-22), a role of trimeric G proteins in the canonical pathway has been difficult to establish and is most likely modulatory (23). In vertebrates, Gαi has been shown to be required in certain contexts of non-canonical Smo signaling including Ca^++^ spiking in the developing spinal cord (24), proliferation of cerebellar granular neurons (25), and activation of the Warburg effect in adipocytes (26).

Here we examine interplay between AKH and Hh signaling in the *D. melanogaster* response to fasting. In response to starvation, Ca^++^ pulses are initiated in the fat body lobes on either side of the brain and propagate along narrow fat body connections to the main fat body lobes along the length of the animal. Signaling from the CC AKH expressing neurons is required for initiation of Ca^++^ pulses in the fat body. In the absence of CC AKH expressing neurons, intracellular Ca^++^ levels in the fat body are low. Fat body Ca^++^ pulses also require Hh signaling. When Hh signaling is blocked in either the fat body or midgut EC, fat body intracellular Ca^++^ levels are elevated and the Ca^++^ pulses are lost. These studies show how signaling from the brain and the midgut and within the fat body are integrated to regulate a key intracellular second messenger.

## Results

### Fasting of early third instar larvae leads to spikes of intracellular Ca^++^ which propagate along the length of the fat body

The fat body of the third instar *D. melanogaster* larva has a complex morphology (Fig 1). There are two fat body lobes in the head region of the larva on either side of the brain and just anterior to the cell bodies of the CC AKH producing neurons. These lobes are linked to the trunk fat body by narrow connections. To examine intracellular Ca^++^ levels in the *D. melanogaster* fat body of early third instar larvae, the GFP Ca^++^ indicator GCaMP6S was employed (27). A *UAS-GCaMP6S* construct was driven in the fat body with *CG-Gal4* (16, 28). In fed larvae, fat body intracellular Ca^++^ remain relatively low and constant (Video 1). In larvae that have been starved for 24 hours, pulses of intracellular Ca^++^ initiate in two lobes of the fat body on either side of the brain and propagate through a narrow fat body connection to the fat body lobes that run along the length of the larvae (Video 2). Pulses advance as a coherent wave (Fig 2A). Pulses of Ca^++^ can also occasionally back propagate from the trunk fat body into the head lobes. The speed of pulse progression varied from 9 to 19 μm/sec, which is within the expected range for intercellular calcium waves (29). With increasing lengths of starvation, the pulses become more robust with increased amplitude (Fig 2 B-E). As an alternative means of assaying intracellular Ca^++^ in the fat body, we generated a construct in which the fat body specific *apolpp* promoter/enhancer was used to directly express GCaMP6S (S1 Fig). This construct was particularly useful when assaying intracellular Ca^++^ in the fat body while manipulating gene expression in a different tissue.

**Fig 1.**
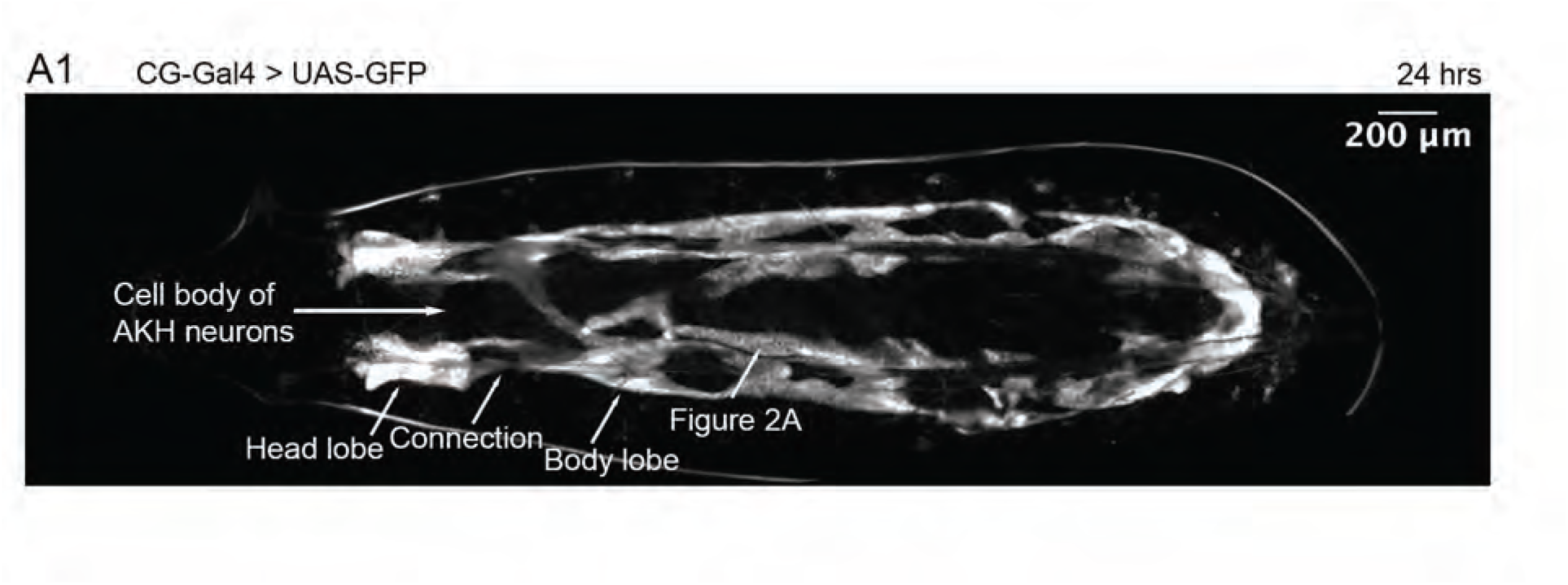
GFP labeling of *D. melanogaster* larva fat body. Early third instar male *w*; CG-Gal4 (7011)/+; UAS-GFP / + larva starved 24 hours to remove gut autofluorescence. The fat body is marked with GFP. Labeled, but not shown, are the cell bodies of the AKH neurons. Also labeled are the fat body lobes in the head, which is linked to the trunk fat body via narrow connections. High resolution imaging of fat body Ca^++^ waves (Fig 2A) was conducted in the fat body region labeled Fig 2A.

**Fig 2.**
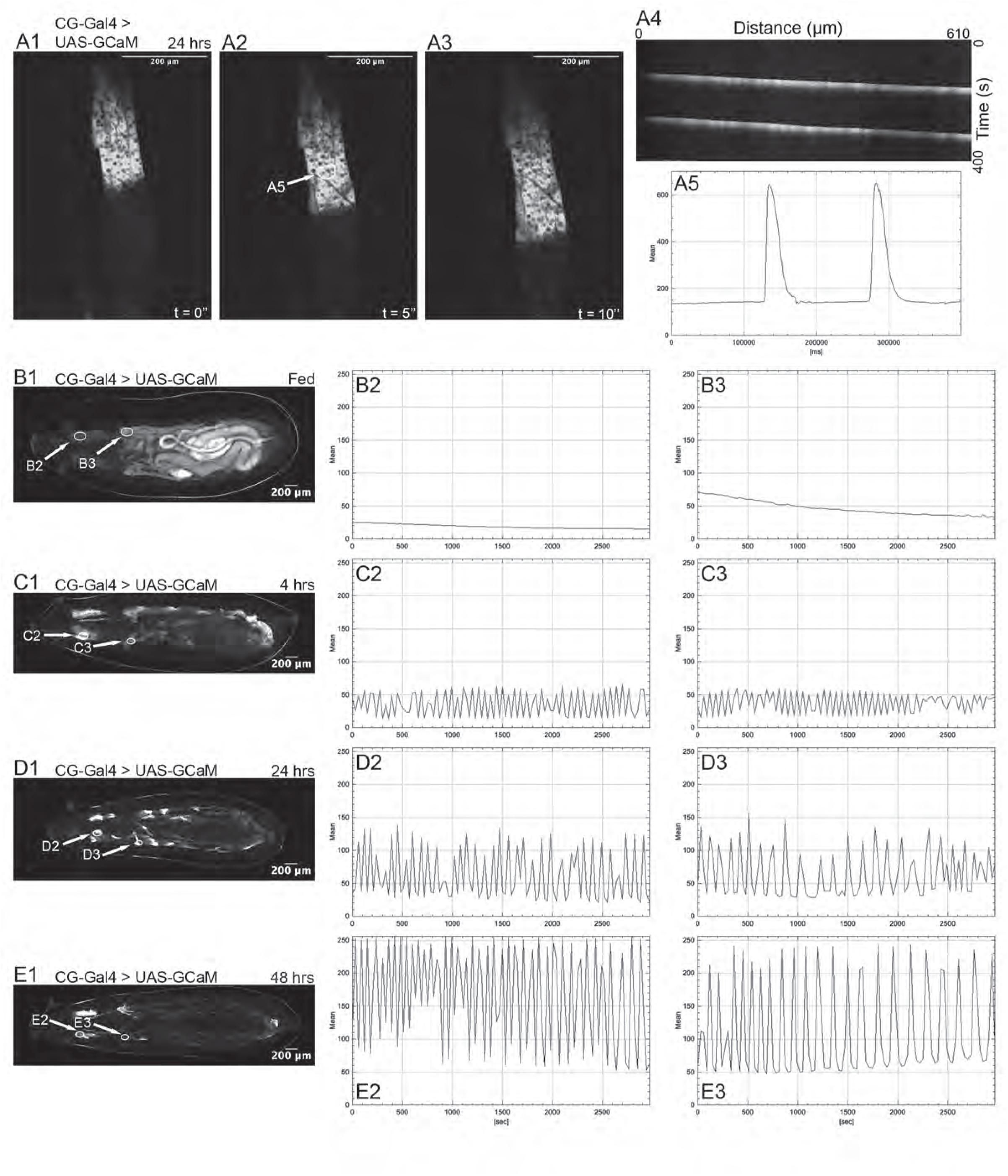
In response to starvation, Ca^++^ pulses are initiated in the fat body lobes on either side of the brain and propagate into the main fat body lobes. All larvae in this figure are *w*; *CG-Gal4* (7011), *UAS-GCaMP6S* (42746)/ + males, and panels B-E were imaged on a wide field fluorescent microscope using identical settings. In this and subsequent figures, the fluorescence intensities are in arbitrary units. (A) High resolution spinning disk confocal imaging of *D. melanogaster* larvae starved 24 hours (time interval 5 sec.) (anterior up). (A1) Ca^++^ levels in fat body visualized at arbitrary time t = 0 second. (A2) Ca^++^ levels in fat body visualized at t = 5 seconds. (A3) Ca^++^ levels in fat body visualized at t = 10 seconds. (A4) Kymograph of fat body over 400 seconds. (A5) GCaMP6S fluorescence levels vs. time. (B) *D. melanogaster* larvae in fed state. (n=9) (C) *D. melanogaster* larvae starved 4 hours. (n=6) (D) *D. melanogaster* larvae starved 24 hours. (n=11) (E) *D. melanogaster* larvae starved 48 hours. (n=11) (B,C,D,E-1) Visualization of Ca^++^ levels in fat body. (B,C,D,E-2) GCaMP6S fluorescence levels vs. time for the marked region of the fat body lobe in the head. (B,C,D,E-3) GCaMP6S fluorescence levels vs. time for the marked region of the medial fat body. It should be noted that there is significant green autofluorescence from the gut in B that disappears following starvation.

**Video 1**. **Live imaging of fed early third instar larva**. *yw*; *UAS-lacZ*/*CG-Gal4, UAS-GCaMP6S* early third instar larva that has been fed. There is significant autofluorescence in the gut with minimal Ca^++^ activity in the fat body.

**Video 2. Live imaging of an early third instar larva that has been starved for 24 hours.** *yw*; *UAS-lacZ*/*CG-Gal4, UAS-GCaMP6S* early third instar larva that has been starved for 24 hours. Strong Ca^++^ pulses are observed along the length of the fat body.

### Signaling from the AKH neurons through AKHR is required for the generation of Ca^++^ pulses in the fat body

Previous work had shown that release of AKH from CC neurons and activation of AKHR in the fat body regulate intracellular Ca^++^ levels and the mobilization of lipids (4-6, 30). We therefore decided to examine the role of AKH and its receptor in the generation of fat body Ca^++^ pulses. AKH signaling was disrupted using two approaches. In the first, the CC AKH expressing neurons were ablated using expression of the cell death inducing gene *rpr* (31). Loss of the AKH expressing neurons removes Ca^++^ pulses in the fat body of larvae that have been starved for 24 hours and leaves fat body intracellular Ca^++^ at low levels (Fig 3A). In the second, fat body Ca^++^ pulses were examined in *AKHR* null mutants. As seen in Fig 3B, loss of *AKHR* function results in the absence of sustained Ca^++^ pulses that originate from the fat body lobes in the head. In half the animals, we unexpectedly observed low frequency fat body Ca^++^ pulses originating from the posterior end of the larva that were not normally present in wild type larvae (Fig 3C). These results demonstrate that the CC AKH expressing neurons and AKHR are required for Ca^++^ pulse generation in the *D. melanogaster* fat body following starvation.

**Fig 3.**
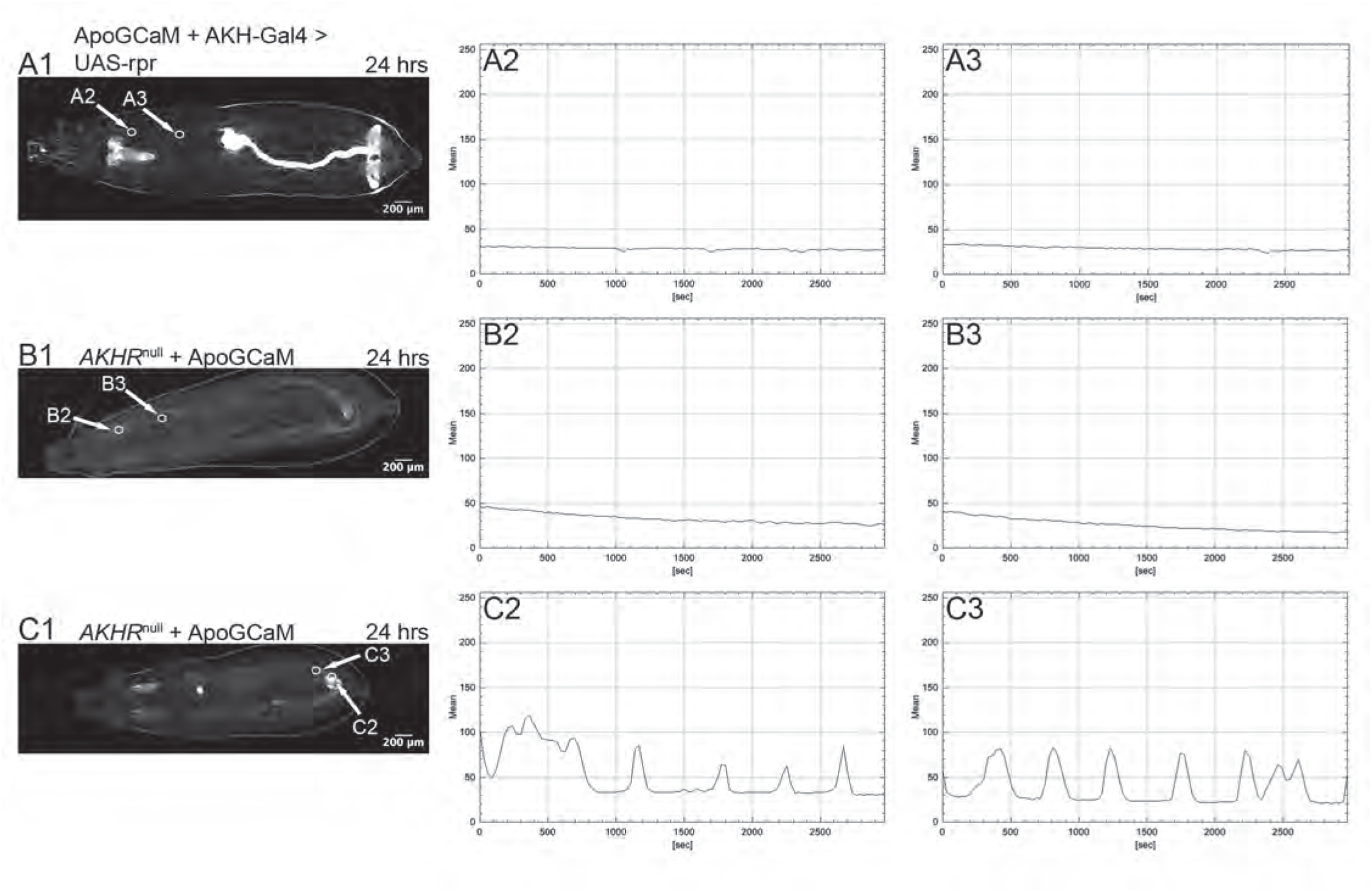
Signaling from the AKH neurons through the AKHR is required for the generation of Ca^++^ pulses in the fat body. (A) *AKH-Gal4* (25683)/*UAS-rpr* (5824); *apolpp::GCaMP6S*/+ larvae identified by the absence of *dfd-YFP* marked balancer chromosomes. (n=6) (A1) GCaMP6S fluorescence levels were visualized in the fat body following 24 hours of starvation. (A2) GCaMP6S fluorescence levels vs. time for the marked region of the fat body lobe in the head. (A3) GCaMP6S fluorescence levels vs. time for the marked region of the medial fat body. The *AKH-Gal4* insertion line has independent expression of GFP in the annal pads, hindgut and nervous system. (B) Homozygous *AKHR*^null^ (80937) females were crossed to *yw hs-flp*; *AKHR*^null^/CyO, *dfd-YFP*; *apolpp:GCaMP6S/*TM6, *dfd-YFP* males. Homozygous *AKHR*^null^ mutants containing *apolpp:GCaMP6S* were identified by the absence of *dfd-YFP* marked balancer chromosomes. (n=6) (C) Same genotype as B. An example of a second class of *AKHR*^null^ larvae in which low frequency Ca^++^ pulses originate in the fat body at the posterior end of the larva. (n=6) (B,C-1) Ca^++^ levels in the fat body visualized after 24 hours of starvation. (B,C-2) GCaMP6S fluorescence levels vs. time for a marked region of fat body lobe in the head. (B,C-3) GCaMP6S fluorescence levels vs. time for a marked region of fat body marked in medial region of larvae

### Knockdown of AKHR effectors disrupt Ca^++^ pulses

Previous work has shown that Gαq, Gγ1, PLC21C and IP_3_R (9) are all required for elevated Ca^++^ levels in the fat body following starvation. To test whether this signaling cascade is also required for Ca^++^ pulses, *UAS-RNAi* constructs targeting each of these components were studied in larvae that had been starved for 24 hours. Knockdown of *G*α*q* with two different RNAi lines resulted in sporadic Ca^++^ pulses in the fat body lobes in the head and poor propagation of Ca^++^ pulses into the trunk fat body (Fig 4A). Knockdown of *Gγ1* resulted in two distinct phenotypic classes. In the first, larvae exhibited low frequency Ca^++^ pulsing in the fat body lobes in the head and slow propagation of Ca^++^ waves into the trunk fat body (n=7) (Fig 4B), and in the second, larvae had elevated Ca^++^ in the fat body lobes within the head region, modest levels of Ca^++^ in the fat body of the trunk region of the larvae, and a lack of pulsing throughout the entire larvae (n=4). Knockdown of *PLC21C* resulted in variable pulsing in the head region and poor pulse propagation into the trunk fat body (Fig 4C). Knockdown of the IP_3_R resulted in a loss of Ca^++^ pulses and low levels of Ca^++^ throughout the fat body (Fig 4D). The low Ca^++^ levels in the knockdown mutants of *G*α*q* and *IP_3_R* and the lack of robust propagation of Ca^++^ pulses in the knockdown mutants of *Gγ1* and *PLC21C* suggest that all of these components function downstream of AKHR to mediate Ca^++^ pulse generation. As it has been shown in *M. sexta* that AKH signaling also regulates Gαs and PKA (10, 11), RNAi was used to target *Gαs*. In larvae starved for 24 hours, knockdown of *Gαs* resulted in a range of phenotypes: three larvae had low frequency Ca^++^ pulsing in the head and intermediate levels of intracellular Ca^++^ in the medial fat body, three lacked Ca^++^ pulsing, had high levels of intracellular Ca^++^ in the fat body lobes in the head, and intermediate levels of intracellular Ca^++^ in the medial fat body, and one larva had robust Ca^++^ waves originating from the posterior end of the larva.

**Fig 4.**
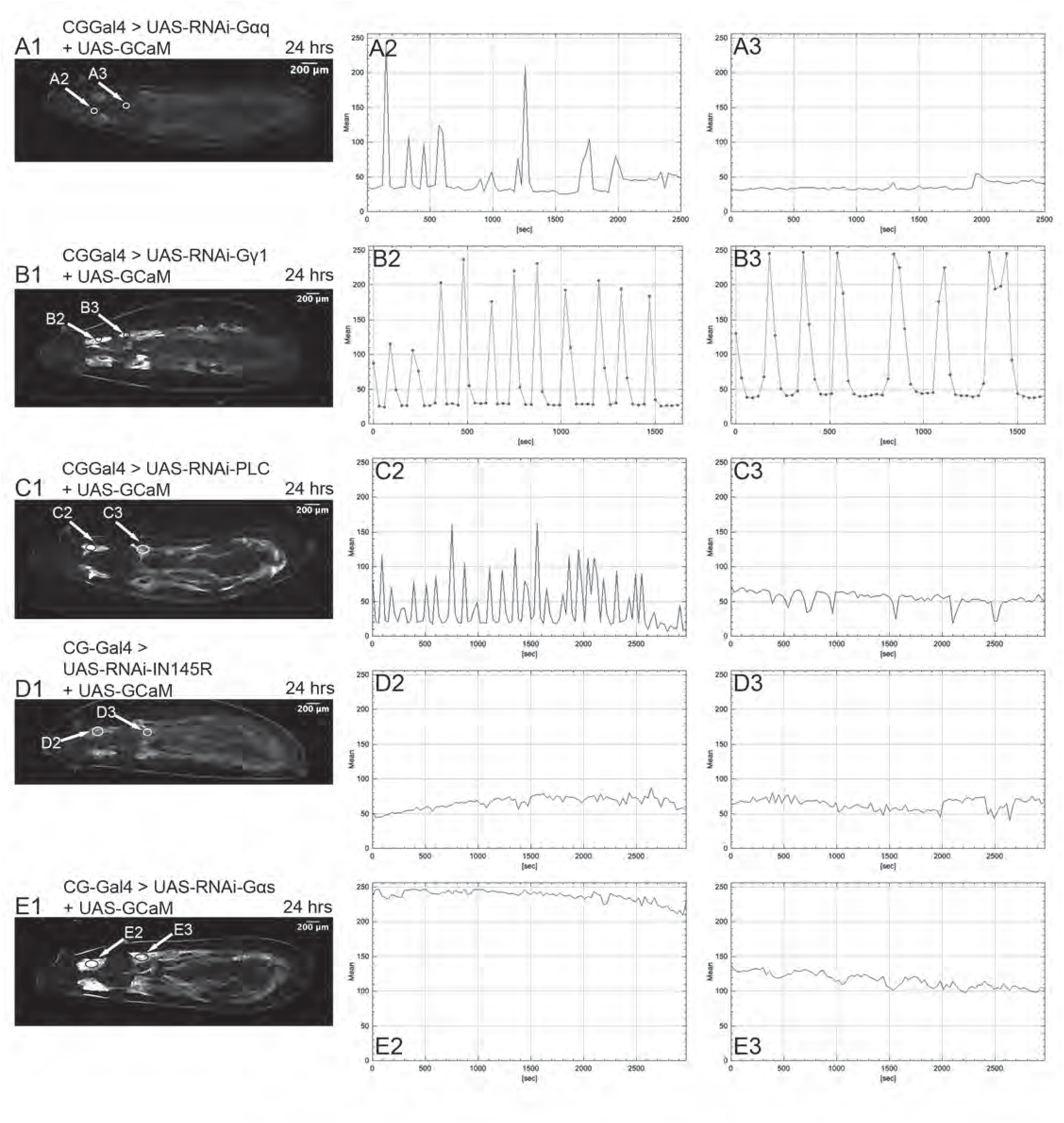
The Gαq, Gγ1, PLC21C and IP_3_R signaling cascade and Gαs are required for fat body Ca^++^ pulsing following starvation. (A) Male *CG-Gal4, UAS-GCaMP6S*/+; *UAS-RNAi G*α*q* (63987)/+ larva starved for 24 hours. (n=6) (B) Male *CG-Gal4, UAS-GCaMP6S*/+; *UAS-RNAi Gγ1* (34372)/+ larva starved for 24 hours. (n=11) (C) Male *CG-Gal4, UAS-GCaMP6S*/+; *UAS-RNAi PLC21C* (32169)/+ larva starved for 24 hours. (n=7) (D) *CG-Gal4 UAS-GCaMP6S*/+; *UAS-RNAi-IP_3_R* (25937)/+ larva starved for 24 hours. (n=11)(All eleven had low levels of Ca^++^, but in one animal a single pulse of elevated Ca^++^ occurred simultaneously throughout the fat body that slowly faded away) (E) *CG-Gal4 UAS-GCaMP6S*/ +; *UAS-RNAi-*Gαs (50704)/+ larva starved for 24 hours. (n=7) (A,B,C,D,E-1) Visualization of Ca^++^ levels in the fat body. (A,B,C,D,E-2) GCaMP6S fluorescence levels vs. time for marked region of a fat body lobe in the head. (A,B,C,D,E-3) GCaMP6S fluorescence levels vs. time for marked region of the fat body marked in the medial region of larvae.

### Loss of Hh signaling leads to elevated levels of Ca^++^ in the fat body and blocks pulse generation

Given the previous studies showing a role for Hh signaling in the release of lipids from the fat body following starvation (16-18), early third instar larvae mutant or knocked down for various components of the Hh pathway were starved for the 24 hours and assayed for fat body Ca^++^ pulsing. In the canonical Hh pathway the Fused (Fu) kinase is required for the full activation of Smo (32) and release of the Ci transcription factor (33). The embryonic development of *fu* mutants can be maternally rescued by heterozygous mothers. As the larvae grow, the maternal contribution of Fu^+^ product is progressively diluted giving rise to variable mutant phenotypes in larval tissues such as the imaginal discs. Here, roughly one half of the *fu*^KO^ (34) larvae that have been starved for 24 hours exhibit loss of Ca^++^ pulsing and elevated Ca^++^ levels throughout in the fat body (Fig 5A). A second approach to inhibit Smo activity made use of the Ptc^Δloop2^ protein, which is unable to bind Hh and constitutively blocks Smo activation (35). Larvae expressing *UAS-ptc*^ι1loop2^ in the fat body consistently exhibited a starvation phenotype like that of the *fu*^KO^ larvae with modestly elevated Ca^++^ levels and a lack of pulsing (Fig 5B). During starvation Hh release from the midgut ECs was shown to be necessary for the activation of Hh signaling in the fat body. Using the *Myo31D-Gal4* driver, we knocked down the expression of *hh* in the midgut ECs, which led to a lack of pulsing in the fat body of starved larvae (Fig 5C). Hh has also been shown to be expressed at low levels in the fat body (18). When RNA was used to knockdown of *hh* in the fat body, Ca^++^ pulsing was also abolished (Fig 5D). Release of Hh is known to require the presence of the Dispatched (Disp) protein in the signaling cells, and its trafficking is thought to involve the Shifted (Shf) protein (36, 37) (*D. melanogaster* homologue of SCUBE2) (38). The *Myo31D-Gal4* driver was used to knockdown *disp* in the midgut enterocytes and results in the elimination of Ca^++^ pulsing in the fat body of starved larvae (S2A Fig). *D. melanogaster shf*^2^ mutants are homozygous viable and fertile with a modest Hh phenotype in the wing. Starved larvae mutant for *shf*^2^ exhibited elevated Ca^++^ levels and a loss of pulsing (Fig 5E).

**Fig 5.**
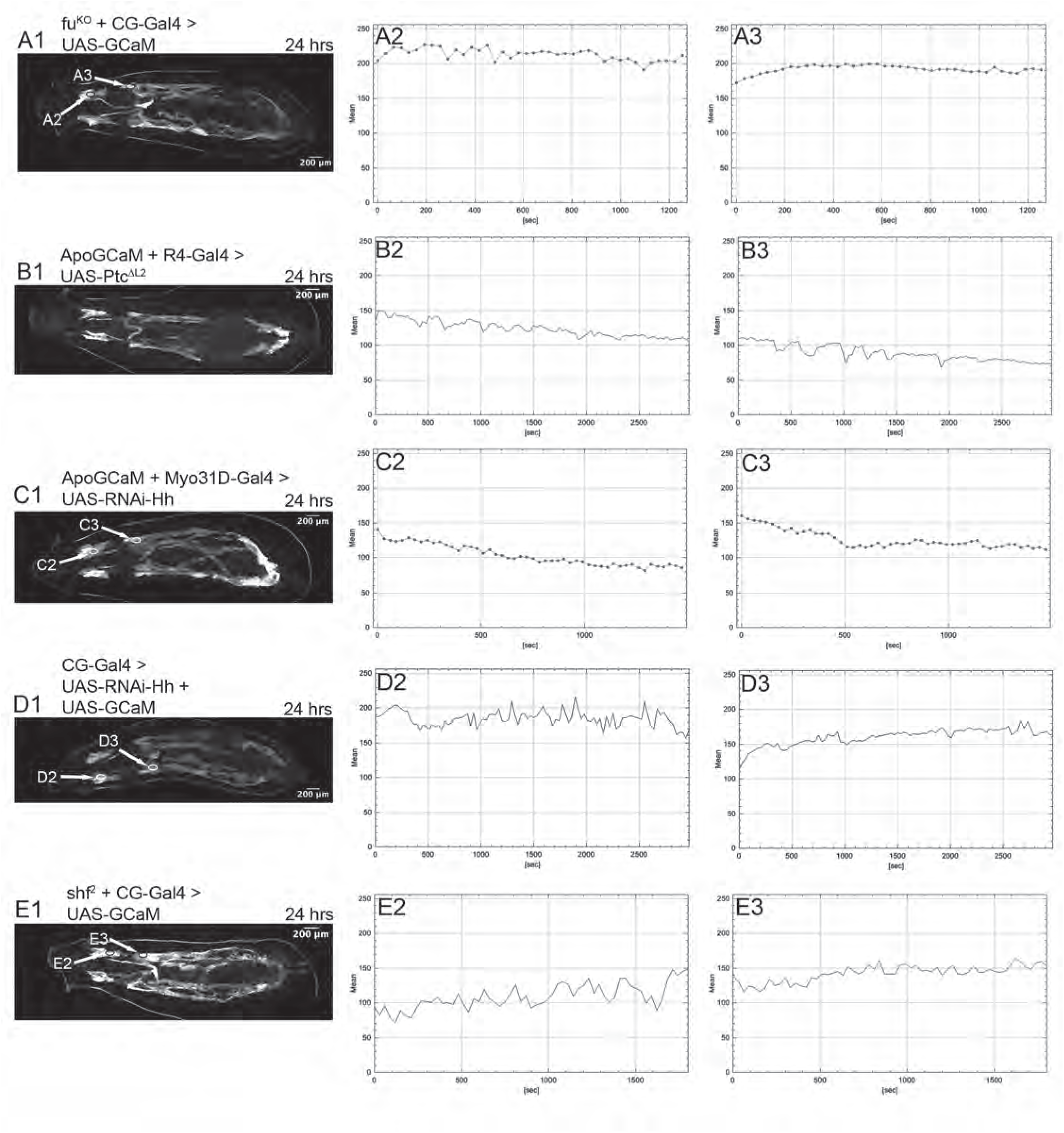
Loss of Hh signaling leads to elevated levels of Ca^++^ in the fat body and blocks pulse generation. Male early third instar larvae were starved for 24 hours. (A) *fu*^KO^ /Y; *CG-Gal4, UAS-GCaMP6S* / +. (n =15 with 7 larvae showing elevated Ca^++^ levels and no pulsing and 8 larvae showing relatively normal pulsing) (B) *UAS-ptc*^Δloop2^/+; *R4-Gal4* (33832)*, apolpp::GCaMP6S* / +. (n=5) (*R4-GAL4* is expressed at higher levels than *CG-Gal4* in third instar larvae.). (C) *Myo31D-Gal4 (P{GawB}Myo31DF*^NP0001^)/ +; *UAS-RNAi-hh* (31042)/ *apolpp::GCaMP6S*. (n =5) (D) *CG-Gal4, UAS-GCaMP6S*/+; *UAS-RNAi-hh* (31042)/+. (n=5) (E*) Shf^2^/Y; CG-Gal4, UAS-GCaMP6S*/+. (n=15) (A,B,C,D,E-1) Visualization of Ca^++^ levels in fat body. (A,B,C,D,E-2) GCaMP6S fluorescence levels vs. time for the marked region of fat body lobe in the head. (A,B,C,D,E-3) GCaMP6S fluorescence levels vs. time for fat body marked in medial region of larvae.

These results suggest that there are two required sources of Hh for Ca^++^ pulse generation, an autocrine signal from the fat body itself and a paracrine/hormonal signal from the mid gut ECs. The downstream components Fu and Ptc are also required along with the Hh chaperones Disp and Shf.

### Generation of Ca^++^ pulses following starvation requires the function of both the transcription factor Ci and G**α**i

Previous work has shown that in response to starvation Hh signaling activates the Ci transcription factor to regulate the expression of *bmm* (18). In the absence of Hh signaling, the Ci protein is processed into a repressor that blocks the expression of Hh target genes (39). Fat body expression of a truncated form of Ci that mimics the Ci repressor resulted in elevated Ca^++^ levels and a loss of pulsing (Fig 6A). As it has been shown in vertebrates that Smo can also couple to Gαi and regulate both intracellular Ca^++^ and metabolism (20-22), animals homozygous for a deletion in the *G*α*i* gene, *G*α*i*^5.4^, that removes the start of translation were analyzed. As can be seen in Fig 6B, loss of *G*α*i* leads to elevation of Ca^++^ levels particularly in the fat body lobes in the head and variable pulsing. Expression of an activated form of Gαi (Gαi^Act^) decreased the frequency of Ca^++^ pulses (Fig 6C). Gαi inhibits Adenylyl Cyclase activity and PKA. Overexpression of the catalytic subunit of PKA, PKA-C1, was used to constitutively activate its function and examine whether the phenotype mimics loss of Hh signaling. Seven animals had varying degrees of pulsing while in three animals, pulsing was lost with moderate Ca^++^ levels (Fig 6D). The requirement for both Ci and Gαi suggests that Hh signaling may act in different ways to regulate starvation induced Ca^++^ pulsing in the fat body. While regulated PKA activity is clearly required for Ca^++^ pulsing in the fat body, its regulation is likely complicated by multiple inputs.

**Fig 6.**
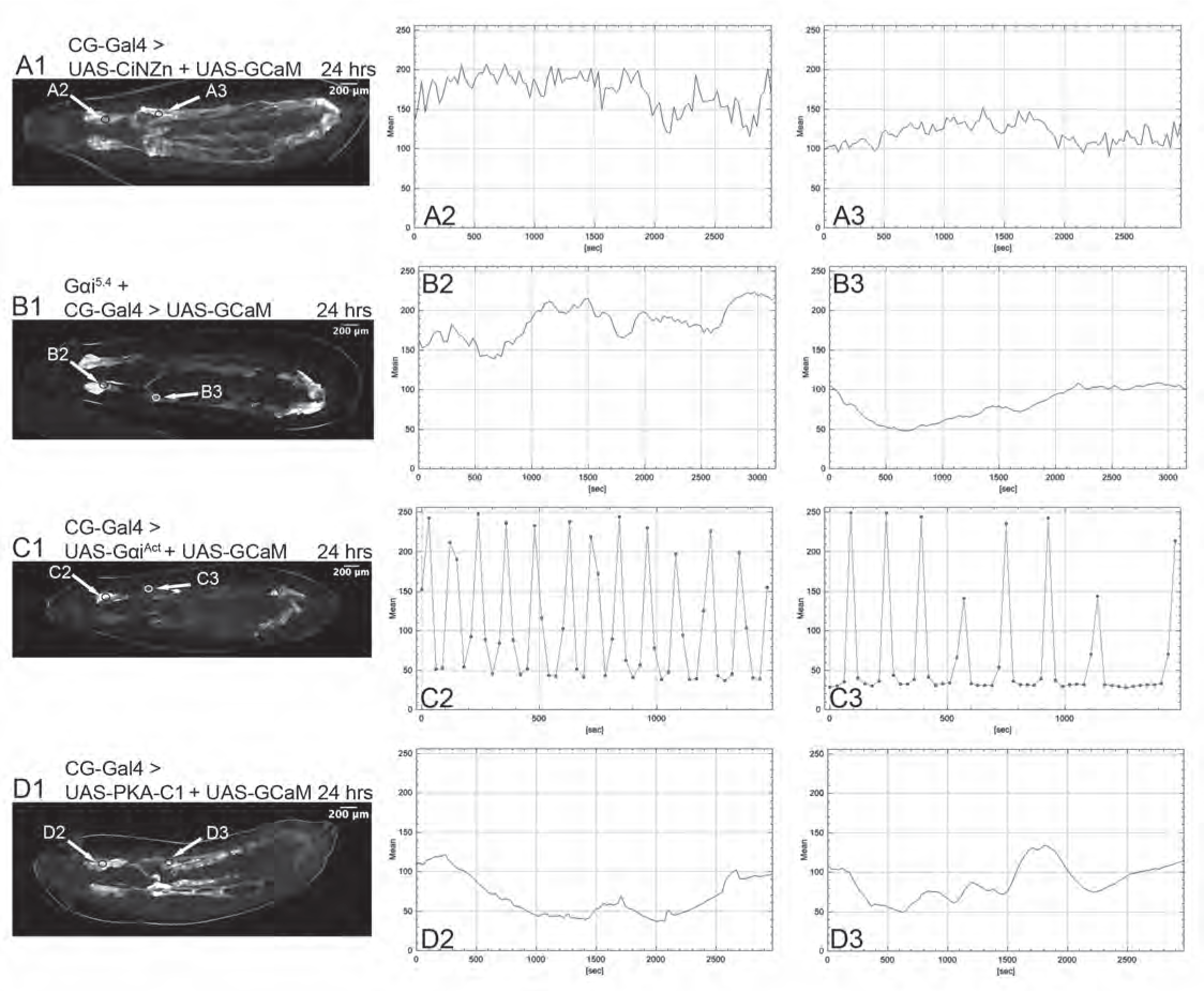
Blocking Ci activity and loss of *G*α*i* inhibit Ca^++^ pulsing. (A) *CG-Gal4, UAS-GCaMP6S*/+*; <u>UAS-ciN[HA]/Zn</u>*/+ (40) male larva. (n=6) (B) *CG-Gal4*/*UAS-GCaPM6S*; *G*α*i*^5.4^/*G*α*i*^5.4^ (n= 6) (C) *CG-Gal4, UAS-GCaMP6S*/*UAS-G*α*i*^Act^ male larva. (n=12) Ten larvae showed low frequency pulsing as shown and two larvae had modest intracellular Ca^++^ levels without pulsing. (D) *CG-Gal4, UAS-GCaMP6S*/*UAS-PKA-C1* (35554). (n=10) (A,B,C,D-1) Visualization of Ca^++^ levels in fat body. (A,B,C,D-2) GCaMP6S fluorescence levels vs. time for marked region of the fat body in the head of the larvae. (A,B,C,D-3) GCaMP6S fluorescence levels vs. time for a marked region of the medial fat body

### Ca^++^ pulsing in the fat body of starved larvae requires SERCA mediated Ca^++^ reuptake

Just as release of intracellular Ca^++^ stores by IP_3_R is required for starvation induced Ca^++^ pulsing in the fat body, we tested whether the return of Ca^++^ from the cytoplasm to the endoplasmic reticulum (ER) required the Sarcoendoplasmic Reticulum Calcium ATPase (SERCA) pump. RNAi knockdown of *SERCA* in the fat body of fed larvae has no effect on intracellular Ca^++^ levels, which remain low (Fig 7A). In the case of larvae that have been starved for 24 hours, knockdown of *SERCA* leads to elevated Ca^++^ levels and a loss of pulsing (Fig 7B).

**Fig 7.**
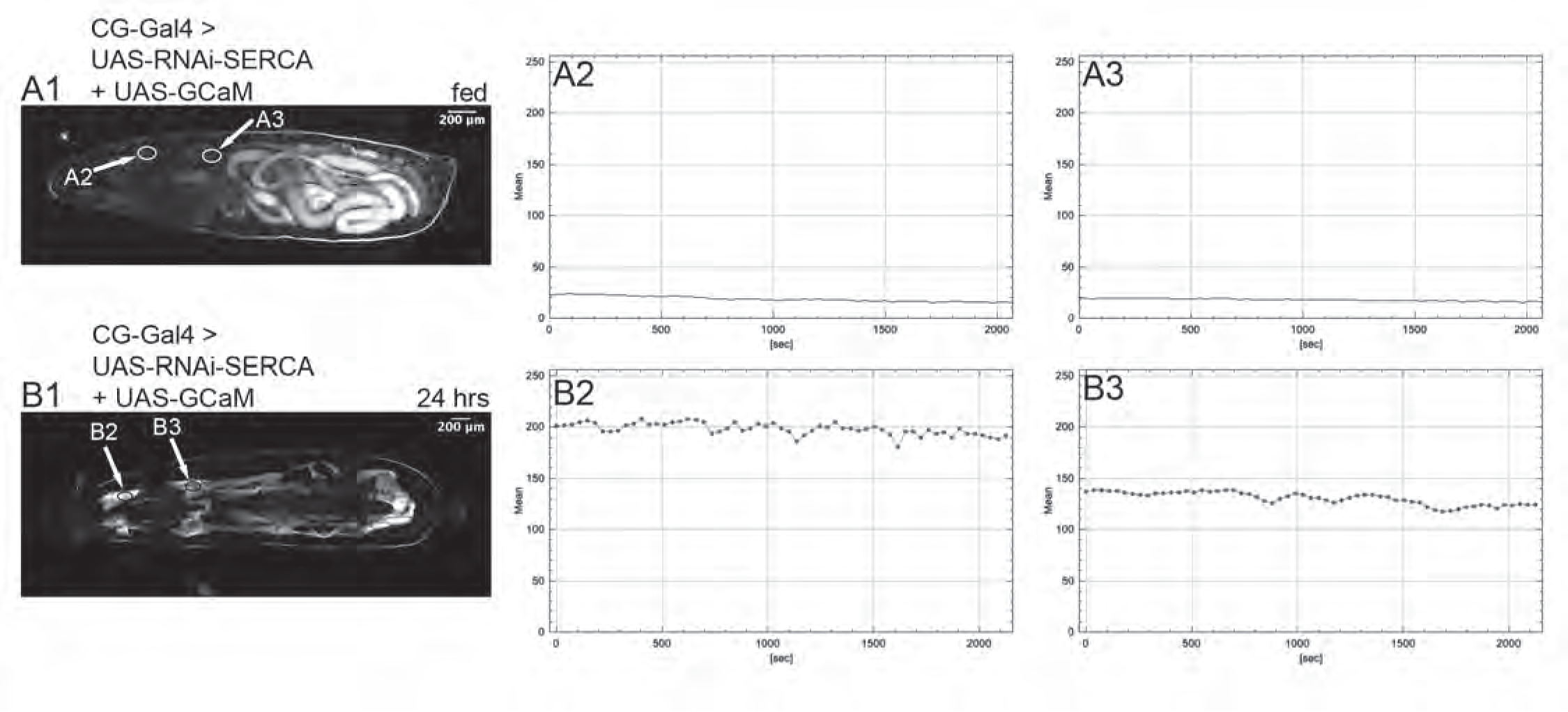
The Sarcoendoplasmic Reticulum Calcium ATPase (SERCA) are required for the generation of Ca^++^ pulses in the fat body. Male *CG-Gal4 UAS-GCaMP6S*/+; *UAS-RNAi-SERCA* (445811)/+ early third instar larvae were fed (n=5) (A) or starved for 24 hours (n=7) (B) and assayed for pulse generation in the fat body. (A,B-1) Ca^++^ levels in fat body. (A,B-2) GCaMP6S fluorescence levels vs. time for marked region in the fat body of the head. (A,B-3) GCaMP6S fluorescence levels vs. time for the marked region of the medial fat body.

## Discussion

### AKH and the AKHR are required for Ca^++^ pulse generation in the fat body of starved larvae

Previous work has shown that ablation of the AKH neurons leads to starvation resistance and reduced trehalose levels in the hemolymph (5) and that a similar phenotype is observed with *AKHR*^null^ mutants (30). AKHR signals through Gαq, Gγ1, and PLC21C to elevate intracellular Ca^++^ levels and mobilize lipid stores in the fat body (9). Experiments in *M. sexta* have shown that AKH can also stimulate the activity of PKA in the fat body and that mobilization of extracellular Ca^++^ leads to lipolysis (10, 11). We have taken advantage of the GFP Ca^++^ indicator GCaMP6S (27) to follow fat body intracellular Ca^++^ levels over time in fed and starved early third instar larvae that have not reached the critical weight for pupation (41). In fed animals intracellular Ca^++^ levels are low and steady. With increasing extents of starvation, Ca^++^ pulses of increasing amplitude are observed in the fat body. These pulses originate in fat body lobes on either side of the larval brain where the CC AKH neurons are located and propagate through a narrow connection to the rest of the fat body. The pulses are dependent on the presence of the CC AKH neurons and AKHR, and in their absence, Ca^++^ levels in the fat body remain low even after starvation. The anatomical division of the fat body into two domains with a narrow linkage suggests that this morphology is somehow used to regulate signaling within the tissue and that the fat body lobes in the head of the larvae act as a pulse generator. Previous work has highlighted the importance of the fat body to the regulation of *D. melanogaster* metabolism; the results presented here show that there is communication within the fat body that is likely important for modulating responses.

In a significant fraction of the starved *AKHR*^null^ mutants, a second source of Ca^++^ pulsing is observed originating from the posterior end of the fat body. It is possible that this second signaling center allows mobilization of fat stores, complements AKH signaling and may explain why *AKHR*^null^ mutants are viable and fertile.

The knockdown of *G*α*q* in the fat body of starved larvae resulted in low frequency Ca^++^ pulses in the fat body lobes in the head and poor propagation of Ca^++^ pulses into the trunk fat body. The phenotype was not as severe as the one observed with the loss of AKHR. This could be a consequence of incomplete knockdown of *G*α*q*, though similar phenotypes were also observed with multiple RNAi lines. It is also possible that the Gβγ subunits have independent roles. In most cases, the knockdown of *Gγ1* resulted in low frequency Ca^++^ pulses as might be expected, but in other larvae, the Ca^++^ levels were unexpectedly elevated. It may be the case that Gγ1 can interact with more than one class of Gα subunit, making it difficult to predict the knockdown phenotype. With knockdown of *PLC21C*, variable pulsing was observed in the head region with poor pulse propagation into the trunk fat body. Again, it may be the case that the RNAi knockdown was not particularly effective with *PLC21C*. However, three different RNAi lines were tried, and all gave similar phenotypes. Mobilization of intracellular Ca^++^ stores is required for pulse generation as RNAi knockdown of IP_3_R blocks their formation. These results show that activation of AKHR, its downstream signaling cascade of Gαq, Gγ1, PLC21C, IP_3_R and release of intracellular Ca^++^ stores are all required for Ca^++^ pulse generation. It is possible that extracellular Ca^++^ also plays a role in pulse generation and/or propagation, as influx of extracellular Ca^++^ is required for the release of lipid stores in vertebrates (42).

### GCaMP6S can act as a Ca^++^ sink and disrupt Ca^++^ signaling

GCaMP6S is a powerful tool for studying intracellular Ca^++^ in living tissue, but to function it must bind Ca^++^. Consequently, GCaMP6S can act as a Ca^++^ sink. Depending on the level of expression, this can disrupt Ca^++^ signaling to varying extents. This was most evident in our experiments using *apolpp::GCaMP6S*. Ca^++^ pulsing was robust in the head lobes of the fat body, but the transmission of the pulses into the trunk fat body was sporadic. We received 10 insertion lines of *apolpp::GCaMP6S* and used the one with the lowest expression level. In all our experiments with *apolpp::GCaMP6S*, only one copy of the transgene was present. In homozygotes with two doses of *apolpp::GCaMP6S*, aberrant high levels and broad waves of intracellular Ca^++^ were regularly observed in the fat body of fed larvae (S1D Fig). We used *CG-Gal4>UAS-GCaMP6S* in most of our experiments to minimize potential Ca^++^ sink artifacts, but it is possible that even this modest expression of *GCaMP6S* sensitizes the tissue, and that in its absence, the intrinsic Ca^++^ pulses would be more robust and extend further into the trunk fat body.

### Hh signaling is required for Ca^++^ pulse generation in the fat body of starved larvae

Previous work has shown that Hh signaling plays a role in various aspects of metabolism in both *D. melanogaster* and vertebrates. Knockdown of Hh signaling leads to elevated levels of triglycerides in the *D. melanogaster* fat body (16, 17, 43), and canonical Hh signaling in the fat body activates the Ci transcription factor increasing the expression of the lipase encoding gene *bmm* (18). In *D. melanogaster* adults, Hh signaling from midgut ECs to the taste sensilla regulates sweet sensation and perception (44). In mouse activation of the Hh pathway in the adipocyte lineage suppresses high fat diet induced obesity (45) and signals through Gαi, extracellular Ca^++^, and Amp kinase to induce a Warburg-like shift in metabolism (26).

Interplay between Hh and Ca^++^ signaling has been identified in a number of tissues. In the *D. melanogaster* lymph gland, Ca^++^ signaling through gap junctions modulates the Hh pathway (46), but in this case the Hh pathway appears to be downstream of Ca^++^ signaling. In *Xenopus laevis* developing spinal cord, Shh increases Ca^++^ spike activity through mobilization of intracellular Ca^++^ stores and Ca^++^ influx (24). Shh can also increase Ca^++^ oscillations in cultured mouse astrocytes (47). In zebrafish mobilization of intracellular Ca^++^ is required for Shh dependent gene expression and appropriate specification of cell fates (48). During the specification of chick feathers, Shh responsive mesenchymal cells have synchronized Ca^++^ oscillations that are disrupted following inhibition of Hh signaling (49). In vertebrates, Hh signaling occurs in the primary cilium. In *X. laevis*, it has been shown that Shh enhances Ca^++^ activity in the primary cilium both through the release of intracellular stores and by influx through the Cation Channel subfamily C member 3 (50).

In the fat body of starved *D. melanogaster* larvae, canonical Hh signaling contributes to the release of lipid stores by upregulating the expression of the *bmm* lipase gene (18). Our results provide strong evidence for second function of Hh signaling to clear cytoplasmic Ca^++^ generated in response to AKH signaling and thereby enable pulse propagation. In the absence of Hh signaling (*fu*^KO^, *ptc*^ΔL2^ and *shf*^2^) in the fat body of starved larvae, Ca^++^ levels are elevated, and pulsing is lost. The phenotype of *G*α*i*^5.4^ mutants was not as severe as other approaches used to disrupt Hh signaling. This suggests there is an additional output from Smo, which may involve direct regulation PKA through PKA binding to a pseudo-substrate site on the Smo C-terminal tail (51). Clearing cytoplasmic Ca^++^ requires SERCA as its knockdown by RNAi leads to elevated Ca^++^ levels and a lack of pulsing in the fat body of starved larvae. In fed larvae, knockdown of SERCA had no effect, suggesting that the elevated Ca^++^ levels in the SERCA knockdown starved larvae were likely dependent on AKH signaling releasing Ca^++^ from intracellular stores. How Hh signaling and the activation of Gαi lead to the clearing of cytoplasmic Ca^++^ is unclear. PKA regulation of SERCA activity has been observed in cardiomyocytes (52) but the sign of the regulation is the opposite of that observed here.

### Ca^++^ pulse initiation and propagation

The Ca^++^ pulses observed in the fat body of starved larvae originate in the head lobes of the fat body on either side of the CC AKH expressing neurons. The simplest explanation for the observed pulse frequency is the possible pulsatile release of AKH from these neurons. These pulses are then propagated through the head fat body lobes and then into the fat body along the trunk of the larvae. It seems likely that the pulses are propagated from cell to cell through gap junctions rather than by paracrine mechanism (29) as the same clusters of cells tend to repeatedly pulse together and the paths of the Ca^++^ pulses seem to follow cells that are tightly connected to each other. The role of Hh signaling in this process appears to be to clear Ca^++^ from the cytoplasm. This is likely to be important for the active regeneration of the Ca^++^ pulse. Active Ca^++^ waves, as opposed to passive ones, are characterized by relatively constant propagation velocity over long distances, which is the case here (29). In the absence of Hh signaling, Ca^++^ levels are high, and pulsing is lost. This may be a consequence of prolonged elevated Ca^++^ inhibiting the function of gap junctions (53). This may explain the difference between the *apolpp::GCaMP6S* starved larvae in S1B Fig vs. S1C In B, the larva has elevated Ca^++^ in the trunk and Ca^++^ pulse propagation to the trunk fat body is lost, while in C, the Ca^++^ levels are low and pulses propagate. Integration of signals to generate coordinated intracellular Ca^++^ waves has also been observed in mouse liver (54).

### Hh signaling and lipid metabolism

The intimate relationship between lipids and Hh signaling has long been noted (55). It has been proposed that the Hh pathway originated as a feedback loop between Ptc and Smo, where Ptc pumped sterols across the plasma membrane and Smo acted as a sensor to down regulate *ptc* expression when sterol levels were sufficiently high. Coupling Hh to Ptc function enabled the pathway to be utilized for intercellular communication and developmental patterning (56). It may be that Smo signaling through Gαi to down regulate PKA activity predates its transcriptional regulation of *ptc*. PKA is an important regulator of metabolism, and its activation leads to lipolysis (57). With the newly available structural data, it has been shown that binding of cholesterol in a deep pocket of Smo appears to necessary for its activation, and Ptc appears to function as a transporter of cholesterol from the inner to the outer plasma membrane (58). Smo sensing of membrane cholesterol could have been used to assess the state of cellular lipids and down regulate lipolysis by down regulating PKA activity. This regulation could become more sophisticated by adding transcriptional regulation of *ptc*, signaling within a tissue by regulation of Ca^++^ levels, and communication between cells through gap junctions. Finally, the addition of Hh regulation of Ptc would integrate communication between tissues, allowing integrated regulation of metabolism and development.

## Materials and Methods

### D. melanogaster strains

The majority of strains used in this study were obtained from the Bloomington Drosophila Stock center. *UAS-ptc^Δloop2^* was obtained from G. Struhl. The *dfd-YFP* balancer chromosomes were obtained from G. Beitel

### *D. melanogaster* transgene construction

*apolpp::GCaMP6S*: The *apolpp* enhancer promoter regions was amplified from a *w^1118^* female fly using NEB Phusion polymerase and the following primers GCC TCG AGC AGT GGT CTC CTG CTG TCA C and GCG GAT CCC ACA CAG ACC ATC CGC GAA TT. It was then cloned as an Xho I, BamH I fragment into corresponding sites of pCaSpeR 4 to generate pCaSpeR-apolpp. The GCaMP6S sequences were amplified from Addgene plasmid #40753 (59) using NEB Phusion polymerase and the following primers GCA GAT CTC GCC ACC ATG GG and CTG ATT ATG ATC TAG AGT CGC GGC CG. The product was digested with Bgl II and Not I and cloned into the BamH I and Not I sites of pCaSpeR-apolpp to generate pCaSpeR-apolpp-GCaMP6S. The splice sites, intron and polyadenylation site from pUAST (60) were amplified using NEB Phusion polymerase and the following primers GGA ATT CGT TAA CAG ATC TTG CGG CC and GC GAA TTC TTG AAT TAG GCC TTC TAG TGG ATC C and cloned into pCaSpeR-apolpp-GCaMP6S using Not I and EcoR I to generate apolpp::GCaMP6S. All PCR inserts were sequenced. Transgenic lines were generated by BestGene Inc.

*UAS-G*αi^act^ was generated from the Gαi cDNA clone (Addgene) LD22201. The Q205L activating mutation (61) was generated by amplifying the Gαi insert as two fragments using NEB Phusion polymerase the primer pairs CCG GAT CCG AAG AGT GCG CGA AGT GAG with CGA TCG C<u>A</u>G GCC ACC CAC ATC GAA AAG TTT G and GGT GGC C<u>T</u>G CGA TCG GAG CG with GGC TCG AGT ACA AAA CCC ACC GGC TGT C. (The underlined bases substitute an L codon for Q.) The two PCR fragments were then used in a second round of amplification with NEB Phusion polymerase and the outside primers CCG GAT CCG AAG AGT GCG CGA AGT GAG and GGC TCG AGT ACA AAA CCC ACC GGC TGT C to generate the Q205L substitution. The fragment was digested with BamH I and Xho I and cloned into the Bgl II, Xho I sites of pUAST to generate UAS-Gαi^act^. The PCR insert was sequenced to confirm the substitution. Transgenic lines were generated by BestGene Inc.

### Generation of *G*α*i* mutation

Imprecise excisions of P{SUPor}Galphai[KG01907] were generated and sequenced. The deletion *G*α*i*^5.4^ starts after nucleotide 19 of the *G*α*i* transcript, inserts the nucleotides TGATGAAATAACATAT and ends in the first intron before the sequence ATGCAACAAGTG removing 1.2 kb, the start of translation, and the first splice donor. This mutation has a similar phenotype to the *G*α*i*^P8^ deletion mutation (62).

### Imaging

Unless otherwise noted, experiments used room temperature (∼22C) third instar larvae (not yet at the critical weight). Animals were reared on molasses medium supplemented with yeast. Larvae were either directly mounted (fed) or placed for the indicated length of time in starvation vials containing 0.75% agar 0.5X PBS. For mounting, a chamber was constructed using two pieces of double-sided Scotch tape. Individual larvae were lightly coated in Halocarbon 400 oil, positioned between the two strips of tape, and immobilized by placing a coverslip on top. Imaging was performed using either a Leica DM6B widefield fluorescent microscope with LED illumination, or in Fig 2A, a Leica spinning disk confocal. All wide field images of GCaMP6S fluorescence were collected using identical settings, and the stitching function of the LAS X software was used to generate composite images. Subsequent analysis was done using ImageJ.

## Abbreviations

CC: corpora cardiaca
EC: enterocytes

## Acknowledgements

We thank Gary Struhl and Greg Beitel for providing *D. melanogaster* stocks. Additional stocks in this study were obtained from the Bloomington Drosophila Stock Center (NIH P40OD018537). Microscopy was performed at the Biological Imaging Facility at Northwestern University (RRID:SCR_017767), MK was the recipient of a Northwestern Summer Undergraduate Research Grant. We would also like to thank Shirley Huang, Nilofar Khanbhai, and Danielle Millan, previous undergraduates in the lab who helped to work out the methodology used in these experiments.

## Supporting information captions

**S1 Fig.**
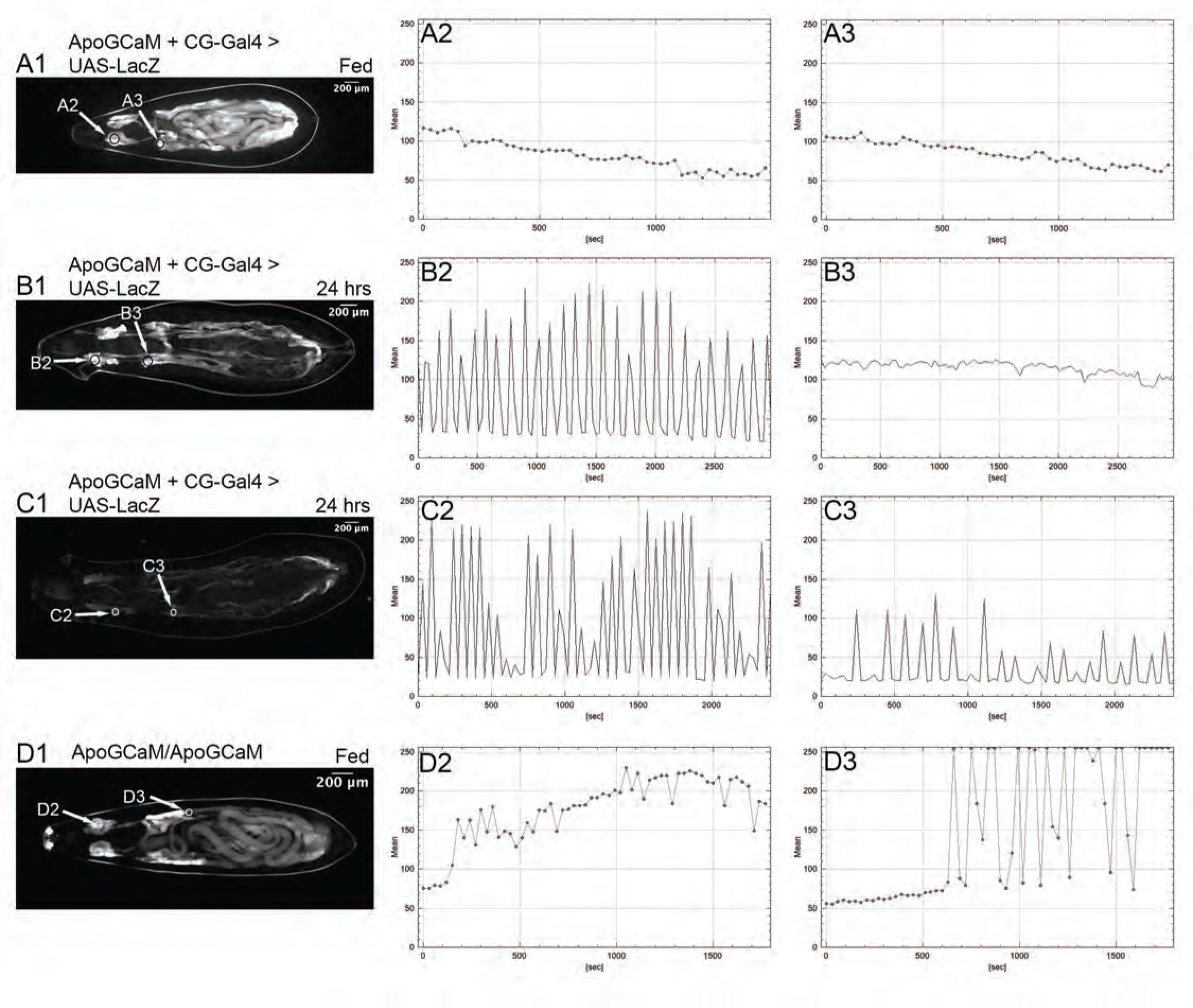
Analysis of Ca^++^ pulses with *apolpp::GCaMP6S*. (A-C) *CG-Gal4*/*UAS-LacZ*; *apolpp::GCaMP6S* male larvae imaged using the same settings as Figure 1. In experiments where *apolpp::GCaMP6S* was used to assay intracellular Ca^++^ in the fat body, there is more variability in the Ca^++^ pulses, and pulse propagation in less robust with roughly half the animals only showing pulsing in the fat body lobes of the head region and the other half having pulsing in the head region and only weak pulse progressing into the fat body lobes of the larval trunk. (A) Fed male larva where Ca^++^ pulses are absent. (B and C) Representative larvae that have been starved for 24 hours. (B) Ca^++^ pulses are only observed in the fat body lobes of the head. (n=20). (C) Ca^++^ pulses from the fat body lobes of the head are only weakly transmitted to fat body lobes of the larval trunk. (n=20) (D) *yw;* CyO*, dfd-YFP/Sp; apolpp::GCaMP6S/apolpp::GCaMP6S* fed larvae (n=6). Even under fed conditions half of the larvae exhibited abnormal, high intensity, Ca^++^ pulses in the larval trunk fat body. In the other half, the larvae exhibited elevated Ca^++^ levels. For this genotype the gain on the camera had to be reduced to avoid saturating the image. This is evident by comparing the gut autofluorescence levels between A1 and D1. (A,B,C,D-1) Ca^++^ levels in fat body. (A,B,C,D-2) GCaMP6S fluorescence levels in a defined region from a fat body lobe in the head region. (A,B,C,D-3) GCaMP6S fluorescence levels vs. time for the marked region of the medial fat body.

**S2 Fig.**
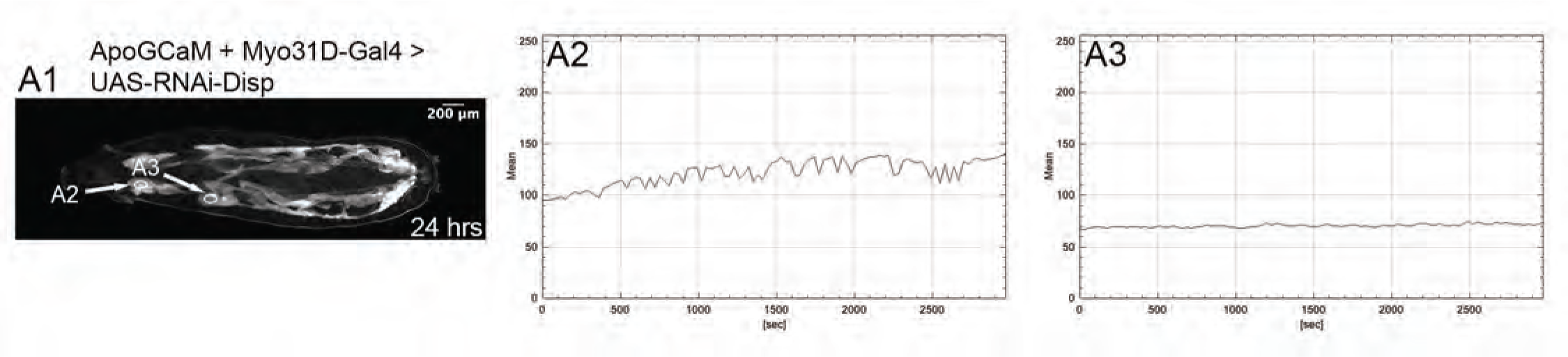
Release of Hh from midgut enterocytes is necessary for the generation of Ca^++^ pulses in starved larvae. (A) *Myo31D-Gal4* / +; *UAS-RNAi-disp* / *apolpp::GCaMP6S* male larva starved for 24 hours. 10 animals had no Ca^++^ pulsing, while two animals had sporatic uncoordinated pulsing in the fat body lobes in the head. (n=12) (A1) Ca^++^ levels in fat body. (A2) GCaMP6S fluorescence levels vs. time for marked region of the fat body in the head of larvae. (A3) GCaMP6S fluorescence levels vs. time for marked region of the medial fat body.

